# Cycad phylogeny predicts host plant use of *Eumaeus* butterflies

**DOI:** 10.1101/2022.12.23.521643

**Authors:** Laura Sierra-Botero, Michael Calonje, Robert K. Robbins, Neil Rosser, Naomi E. Pierce, Cristina López-Gallego, Wendy A. Valencia-Montoya

## Abstract

*Eumaeus* butterflies are obligate herbivores of *Zamia*, the most diverse neotropical genus of cycads. *Eumaeus-Zamia* interactions have been mainly characterized for species distributed in North and Central America. However, host plant use by the southern *Eumaeus* clade remains largely unknown, precluding a comprehensive study of co-evolution between the genera. Here, we combine fieldwork with museum and literature surveys to expand herbivory records for *Eumaeus* from 21 to 38 *Zamia* species. We inferred a time-calibrated phylogeny of *Eumaeus* to test for distinct macroevolutionary scenarios of host plant conservatism and co-evolution. We found remarkable coincidence between *Eumaeus* and *Zamia* diversification, with the butterfly stem group diverging at the same time as the most recent radiation of *Zamia* in the Miocene. Cophylogenetic reconciliation analyses show a strong cophylogenetic signal between cycads and their butterfly herbivores. Bipartite model-based approaches indicate that this is because closely related *Zamia* species are used by the same *Eumaeus* species, suggesting host plant resource tracking by the butterfly herbivores. Our results highlight a case of tight evolution between *Eumaeus* butterflies and cycads, pointing to the generality of correlated evolution and phylogenetic tracking in plant-herbivore interactions across seed plants.

## Introduction

Cycads are tropical gymnosperms and among the oldest living lineages of seed plants, with a fossil record extending back over 265 million years (Zhifeng & Thomas, 1989). Cycads are considered “living fossils” since some morphological traits from species in the Mesozoic are remarkably well preserved in current lineages (Norstog & Nicholls, 1997). Despite these ancient origins, all cycad genera have undergone recent radiations across the tropics (Nagalingum et al., 2011). Little research, however, has focused on ecological interactions between cycads and animals. Such work is crucial not only to understand the evolution of these interactions in plants other than angiosperms, but also to better characterize the population dynamics of cycads and associated species in present-day ecosystems. This is particularly pressing given that cycads are one of the plant taxa with the highest risk of extinction, rendering their obligate pollinators and herbivores similarly imperiled (*IUCN Red List of Threatened Species*, 2022).

As gymnosperms, cycads have commonly been thought to have few ecological interactions with animals. However, this view was challenged by work in the 1980s demonstrating beetle-mediated pollination (Norstog & Fawcett, 1989; Tang, 1987) and specialized insect herbivory (Bowers & Larin, 1989; Clark & Clark, 1991; Rothschild et al., 1986). It is now known that all living cycads engage in obligate and highly specialized interactions with beetles and thrips for pollination (Toon et al., 2020). In addition, butterflies, moths, and beetles are specialist herbivores of many cycads (Salzman et al., 2018). Cycads exhibit a remarkable repertoire of chemical defenses, comprising compounds known to be highly neurotoxic and carcinogenic to some animals (Charlton et al., 1992; Duncan et al., 1988; Laqueur & Spatz, 1968). The best-characterized compounds include B-Methylamino-L-alanine (MAM) and the non-protein amino acid methylazoxymethanol acetate (BMAM), both found in the tissues of all cycad genera (De Luca et al., 1980; Vega & Bell, 1967).

Most cycad-feeding insects are lepidopterans (butterflies and moths), with host plant records from nine genera in the families Erebidae, Cosmopterigidae, Tineidae, Nymphalidae, Blastobasidae, Geometridae, and Lycaenidae (Whitaker & Salzman, 2020). Despite this taxonomic diversity, only lycaenid butterflies are known to have cycads as obligate caterpillar hosts. The genera *Luthrodes* and *Theclinesthes* feed on *Cycas* and *Macrozamia* in Asia and Oceania, respectively (Whitaker & Salzman, 2020). In the Neotropics, *Eumaeus* are the only known obligate herbivores of cycads, feeding on Zamiaceae. Early-divergent species of *Eumaeus* are able to feed on genera such as *Dioon* and *Ceratozamia*, but the majority use *Zamia*, the most diverse neotropical genus of cycads, as their primary host plants.

Since all *Eumaeus* species are obligate herbivores of Zamiaceae, the cycad-feeding habit is inferred to have evolved in the last common ancestor of the genus (Robbins et al., 2021). The shift to feeding on cycads seems to have been accompanied by the rapid evolution of aposematism and conspicuously gregarious, non-ant-associated larvae, together, a unique phenotype within the family Lycaenidae (Pierce et al., 2002a; Robbins et al., 2021). While many lycaenid caterpillars are cryptically colored and form associations with ants (Pierce et al., 2002b), *Eumaeus* caterpillars are generally not ant associated and have aposematic, bright-red coloration with yellow or white bands. The adults of the majority of species also have red elements on the body and wings. Accordingly, toxic BMAM and MAMM compounds sequestered from cycads render all life stages (eggs, larvae, and adults) unpalatable to predators (Castillo-Guevara & Rico-Gray, 2002; Rothschild et al., 1986; Schneider et al., 2002). Caterpillars are able to detoxify these compounds from their host plants, as recent genomic evidence suggests that the rapid evolution of protein families related to autophagy and phagocytosis likely underlies these adaptations (Robbins et al., 2021).

These findings add to a growing number of examples of butterfly detoxification of plant toxins (Edger et al., 2015; Matsubayashi et al., 2010; Wheat et al., 2007; Winkler & Mitter, 2008). Ehrlich and Raven (1964) proposed that the arms race between the evolution of toxic secondary compounds in plants and the associated detoxification mechanisms in butterflies results in a step-wise, reciprocal pattern of co-evolution. The emerging consensus from the majority of plant-insect interaction studies does not support the strict “escape and radiate” interpretation of this model, but nevertheless points to strong phylogenetic conservatism of insect-host plant associations (Allio et al., 2021; Forister et al., 2015). The literature on reciprocal evolution is significantly biased toward interactions with flowering plants, and little is known about macroevolutionary patterns of herbivory in other plant groups. Therefore, the *Zamia* – *Eumaeus* system provides an excellent opportunity to understand the ecology and evolution of highly-specialized herbivory beyond angiosperms and test the generality of this pattern across seed plants.

Although *Eumaeus – Zamia* is an ideal system to test for phylogenetic conservatism of host plant associations, there is a significant gap in natural history knowledge for species occurring in the southern distribution of the range of *Eumaeus*. The three northern species, *Eumaeus childrenae, Eumaeus atala*, and *Eumaeus toxea* are distributed from Florida to Mexico and have been widely collected and studied (see (Contreras-Medina et al., 2003; Jiménez-Pérez et al., 2017; Koi & Daniels, 2015, 2017; Koi & Hall, 2016; Martínez-Lendech et al., 2007; Ruiz– García, 2020). The southern species, *Eumaeus godartii, Eumaeus toxana*, and *Eumaeus minyas* form a monophyletic lineage (Robbins et al., 2021) distributed from Costa Rica to Northern Bolivia, and are hereon referred to as the “southern clade.” In contrast to the northern species, the host plants and geographical distributions of the southern clade are poorly known, and few studies on individual species have been conducted (Castillo-Guevara & Rico-Gray, 2002, 2003; González, 2004; Santos Murgas & Abrego, 2016; Segalla et al., 2021; Taylor B., 2020).

This scarcity of data for the southern clade of *Eumaeus* has thus far precluded a genus-level study of phylogenetic patterns underlying host plant use. Here, we combine fieldwork with a review of collections and literature to fill in gaps in geographical distributions and host plants for species in the southern *Eumaeus* clade, particularly in Colombia. Colombia is not only the world’s most species-rich country for *Zamia* (López-Gallego et al., 2019) and *Eumaeus*, but it is also the country that connects species in the Amazon with those in the Central-American and Caribbean regions, via the Darién and Andean forests. We build the most comprehensive dataset to date on distributions and host plant use for the genus, and use it to test co-evolutionary hypotheses about the *Zamia* - *Eumaeus* interaction.

## Materials and Methods

### Sampling in natural populations

Field studies were carried out in natural populations at key localities of the *Zamia* distribution in Colombia. We explored four localities encompassing distinct biogeographic regions ranging from the Pacific to the Amazon. These were: 1) “Cañón del Río Alicante”, Maceo, Antioquia, in the Magdalena Valley region (6°33’3’’ N, 74°54’3’’ W, 500 masl, January, 2021). 2) “Hacienda La Mejía”, Chigorodó, Antioquia, in the Darién region (7°31’21.” N, 76°35’2”W, 80 masl, June, 2021). 3) The Tropical Forestry Center “Pedro Antonio Pineda”, Buenaventura, Valle del Cauca, in the Pacific region (3°57’1’’ N, 76°59’2’’ W, 60 masl, December 2021). 4) Biological station “El Zafire”, Leticia, Amazonas, in the Amazon (4°0’32.0’’ S, 69°53’4’’ W, 118 masl, in January, 2022).

In each of the localities, adult *Eumaeus* butterflies were collected manually using entomological nets. Specimens were stored in glassine envelopes and deposited in the Biological collection of CES University (CBUCES) and the Entomological collection of the University of Antioquia (CEUA) (Medellin, Colombia) biological collections for later identification. Host plant and geographical coordinates were recorded in the field upon collection.

### Host plant association survey and identification of specimens

We surveyed *Eumaeus* specimens held in the main entomological collections in Colombia, including those at the CES University in Medellín (CBUCES), the University of Antioquia (CEUA), the Francisco Luis Gallego Entomological Museum at the National University of Colombia in Medellín (MEFLG), the Natural History Museum at the National University of Colombia (ICN-L), and the Alexander von Humboldt Biological Resources Research Institute (IAvH-E). We also surveyed museums such as USNM in Washington, US; MUSM in Lima, Peru and BMNH in London, UK.

In addition, we compiled photographic records of *Eumaeus* taken on or near *Zamia* during field trips by the Colombian Cycad Society, an academic NGO dedicated to cycad research and conservation. Most of these photographs were of either the adult or larval stages, meaning we could reliably identify the *Eumaeus* species as well as its host plant. We also checked and compiled records of adult *Eumaeus* from online biodiversity databases (iNaturalist, Butterflies of America, GBIF) and the web pages of natural history museums (the Yale Peabody Museum of Natural History, the Museum of Comparative Zoology of Harvard University, the McGuire Center for Lepidoptera and Biodiversity, Auckland Museum Entomology Collection, Alfonso L. Herrera Zoology Museum in the National Autonomous University of Mexico, the Royal Belgian Institute of Natural Sciences Collection, the National Institute of Biodiversity of Costa Rica and the Museum of Natural History of Mexico City).

All museum and field-collected specimens and the available photographs were identified using Robbins et al. (2021), Goodson (1947), and Butterflies of America (Warren et al., 2016). Key morphological characters included: coloration, size, wing shape, presence and position of wing spots, and presence of tibial spurs. Most specimens were identified based on external morphological characters, but genitalia were dissected for specimens with ambiguous characters or anomalous localities. When host plant data were not available for a record, we inferred the host species based on the known distribution of *Zamia* species, unless multiple *Zamia* species were known to occur at the locality. Using all of the available records, we then mapped the distribution of *Eumaeus* using QGIS 3.26.1 (QGIS Development Team, 2022).

### Inference of phylogenetic associations between Eumaeus and Zamia

We used the data from fieldwork, biological collections, biodiversity platforms and literature to build a presence/absence matrix of host plant use by *Eumaeus*. To test for broad cophylogenetic signal between *Zamia* and *Eumaeus*, we used the R package `paco: Procrustes Application to Cophylogenetic Analysis` (Balbuena et al., 2013). This method is the most recent and widely-used global-fit approach to cophylogeny (Blasco-Costa et al., 2021) `paco` uses the phylogenies of both groups and the interaction network to test whether the observed matrix is more congruent with the evolutionary history of both partners than a random assemblage of these interactions (Balbuena et al., 2013). We used the r0 algorithm, which assumes that *Eumaeus* tracks the phylogeny of *Zamia*. This algorithm is the adequate model for herbivores and parasites, yet to standardize both phylogenies prior to superimposition, the argument “symmetry” in the model was set to TRUE. This means that both phylogenies are standardized prior to superimposition resulting in the best-fit of the superimposition being independent of both phylogenies (Hutchinson et al., 2017). Cophylogenetic signal was considered to be significant when it was smaller than 95% of the values obtained from 1000 randomisations of the aggregated interaction dataset (Fuzessy et al., 2022).

Since `paco` tests for overall congruence of the phylogenies of the interacting groups, but does not partition the contribution of the different phylogenies to the cophylogenetic signal, we additionally fitted linear mixed models. To test for phylogenetic signal in *Zamia*-*Eumaeus* interactions we built a phylogenetic generalized linear mixed model (PGLMM), using the R package phyr (Li et al., 2020). PGLMMs treat the strengths of pairwise interactions (in this case, host plant use by butterflies) as the response variable and incorporate phylogenies as anticipated covariances among these interactions (Rafferty & Ives, 2013). The model structure is identical to equation 3 in Rafferty and Ives (2013), except we specified a binomial error distribution because the response variable Y comprises presence/absence data. The model contains nested phylogenetic covariance matrices describing three patterns. The first describes the pattern in which closely related butterflies are more likely to use the same host plant (covariance matrix *c*). The second describes the pattern in which a butterfly species is more likely to use a set of closely related host plants (covariance matrix *g*). The third describes the pattern in which closely related butterflies are more likely to use closely related host plants (covariance matrix *h*). The significance of these phylogenetic effects can be tested by dropping them and applying likelihood ratio tests (LTRs). The covariance matrices c and g are nested components of covariance matrices h. Therefore, we tested the significance of h by dropping it from a complete model including all terms (c + g + h), but c and g were tested by dropping them from a reduced model which did not contain the other two terms. Of the 36 species of *Zamia* recorded as host plants of *Eumaeus*, 33 were used in the phylogenetic analysis; the remaining species were not included in the available phylogeny Calonje et al., 2019). We scaled the *Eumaeus* tree by branch length and transformed it into an ultrametric topology using the chronos function (lambda=0) in APE (Paradis & Schliep, 2019).

### Estimation of divergences times in the Zamia and Eumaeus phylogenies

As testing for co-evolution entails accounting for the topological history as well as the diversification timing of the interacting groups, we used the most recent time-calibrated molecular phylogeny of *Zamia* (Calonje et al., 2019). For *Eumaeus*, we inferred an ultrametric genome-wide phylogeny using sequence data published by Robbins et al. (2021). We used MCMCtree v4.9 for divergence time estimation, incorporating secondary node calibrations based on a recent fossil-calibrated phylogeny of Lepidoptera and Eumaeini (Espeland et al., 2018; Valencia-Montoya et al., 2021). The remaining node age priors were set to uniform. We performed an approximate likelihood calculation of divergence time in MCMCtree with the estimation of the gradient and Hessian matrix of the branch lengths. We then ran the concatenated alignment under the F84 substitution model and gamma with five rate parameters.

## Results

### Geographic distributions and host plants of Eumaeus

On the southern Pacific coast of Colombia, we found *Eumaeus godartii* feeding on *Zamia chigua* and *Z. amplifolia* (Fig 1k-n), which co-occur near Buenaventura. We also uncovered museum specimens indicating that the *E. godartii* occurs in the Darién region close to the border with Panama (Fig. 2b, yellow dots). *Eumaeus minyas* was collected at Maceo in the central Andean cordillera on the eastern slope of the Magdalena river, where it was using *Z. incognita* as host plant (Fig 1j). This population was previously described in the literature as *E*. cf. *godartii* (Valencia-Montoya et al., 2017), but all the diagnostic characters suggest it is *E. minyas* (following Robbins et al., 2021). On the basis of museum records, we here extend the distribution of *E. minyas* throughout the Magdalena river region in the Colombian Andes (Fig. 2b, blue dots). Notably, specimens exhibiting intermediate morphological traits between *E. minyas* and *E. godartii* were found in transitional zones in the central Andean cordillera, between the Cauca and Magdalena river valleys (Fig. 2b violet dots, SI Appendix). *Eumaeus toxana* was found in white-sand forests near Leticia in the Colombian Amazon region, using *Z. cupatiensis* as host plant (Fig 1a-d).

**Fig. 1.**
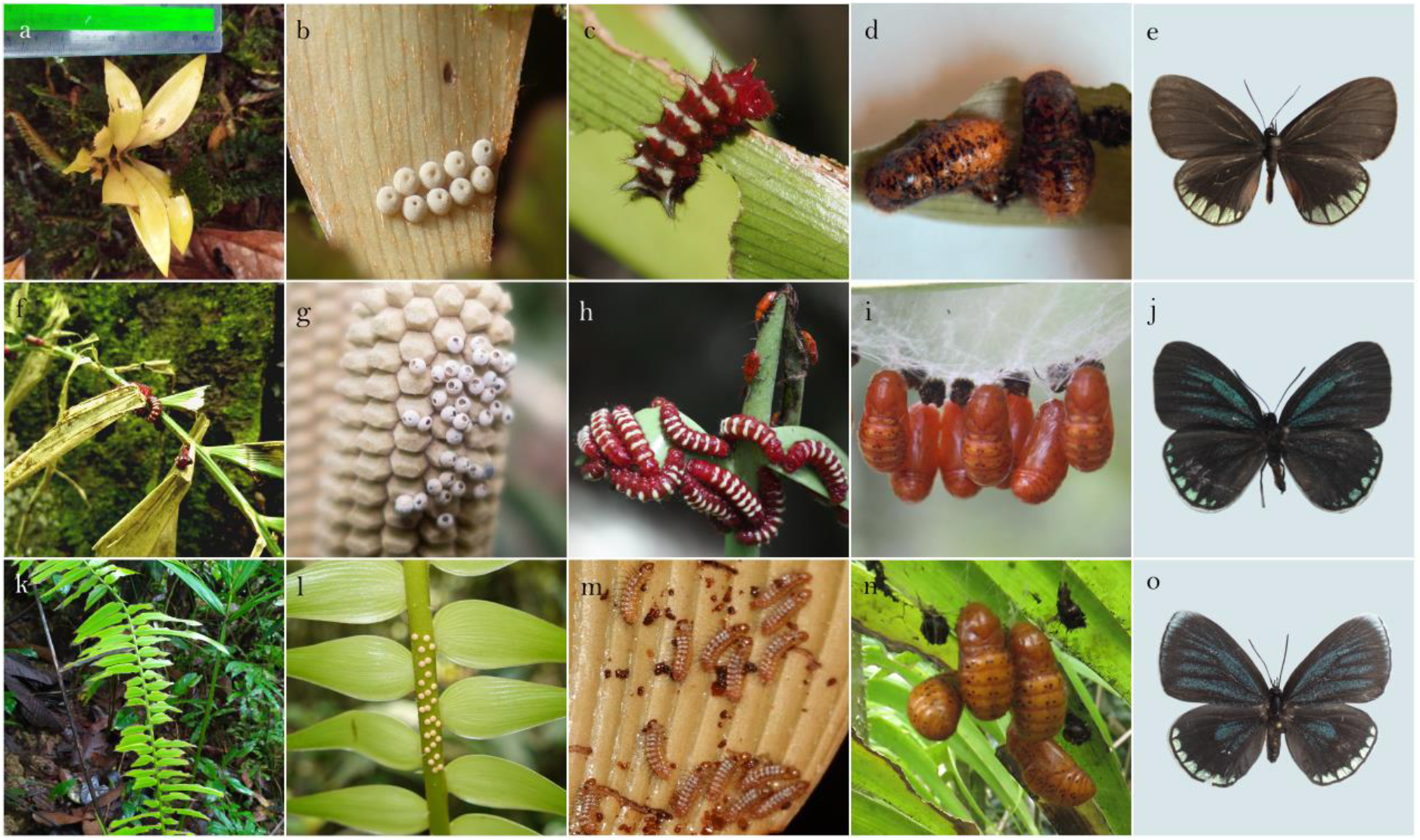
Species of the southern *Eumaeus* clade. As with other *Eumaeus* species, immatures develop only on cycads, specifically *Zamia*. **a-e:** *E. toxana*. **a**. *Z. cupatiensis;* the host plant species of *E. toxana* in the Amazon. **b**. Egg clutch on *Z. cupatiensis* leaf **c**. Larva on *Z. cupatiensis* leaf. **d**. Late stage pupae **e**. Dorsal view of an adult female *E. toxana* from Vaupés. **f-j**: *E. minyas* **f**. Leaf of *Z. tolimensis* affected by herbivory. **g**. Egg clutch on female cone of *Z. incognita*. **h**. Larvae on *Z. encephalartoides*, with another reported cycadivore *Aulacoscelis sp* (Coleoptera: Orsodacnidae) visible in the background. **i**. Pupae on *Z. encephalartoides*. **j**. Dorsal view of female adult from Girón, Santander. **k-o**. *E. godartii*. **k**. *Zamia chigua* leaf affected by herbivory. **l**. Egg clutch on rachis of *Z. chigua*. **m**. Newly emerged larvae on *Z. amplifolia* leaflet. **n**. Pupae hanging on the underside of *Z. chigua* leaf. **o**. Dorsal view of adult female from Parque Natural los Katíos, Riosucio. Credits: Jonathan Castro (g) and Cornelio A. Bota-Sierra (i).

**Fig. 2.**
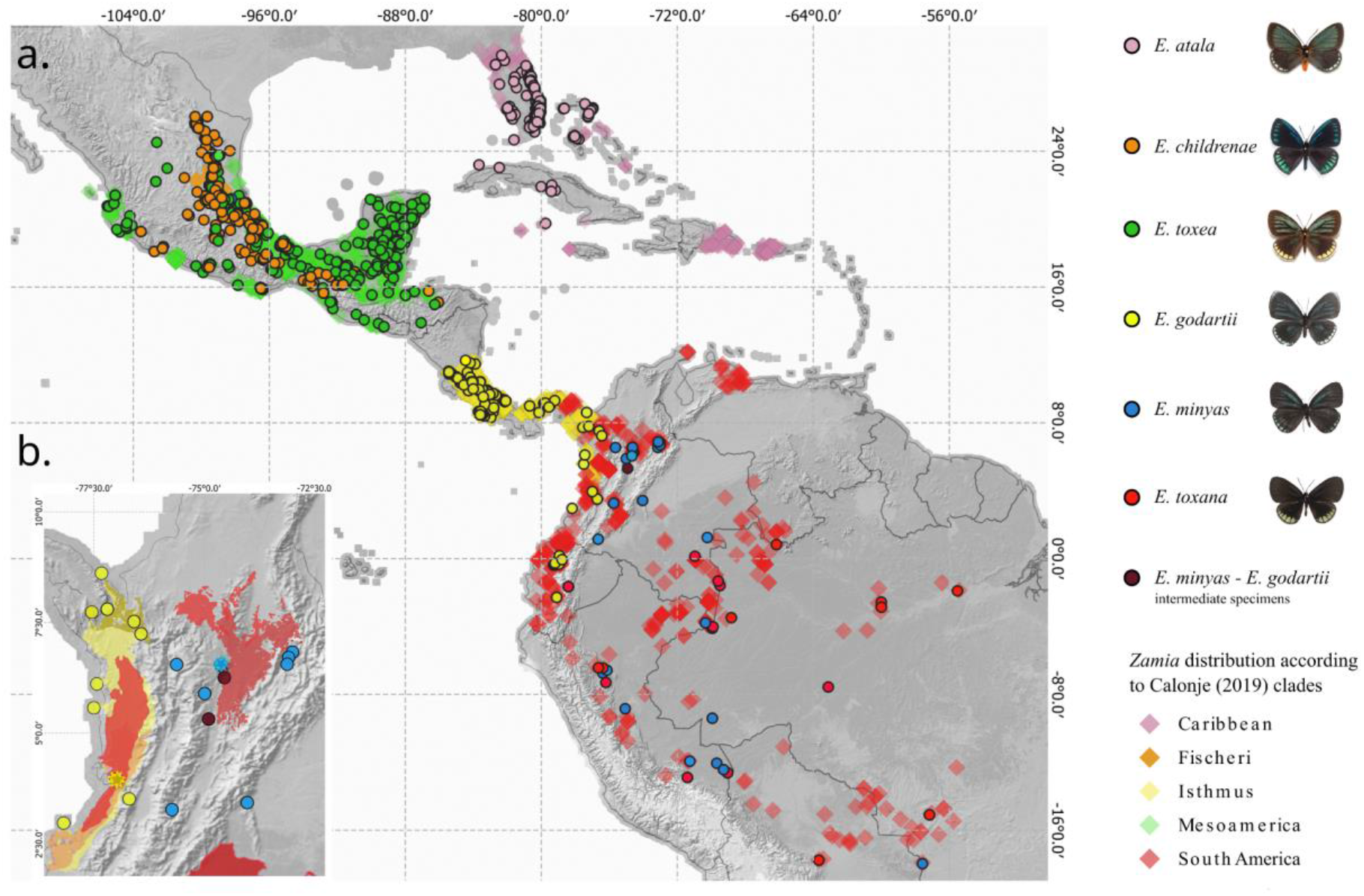
**a**. Revised distribution of *Eumaeus* species in the Neotropics, based on additional fieldwork and specimen surveys. **b**. Magnified view of Colombia showing *Eumaeus* point localities superimposed on *Zamia* range maps. Asterisks within the map show study sites. Credits: Photographs of the northern clade species (*E. atala, E. childrenae, E. toxea*): Butterflies of America.

After reviewing museum specimens and online data sources and coupling them with our field data, we were able to create a significantly improved continental-scale distribution map of *Eumaeus*. We confirmed that *E. atala, E. toxea*, and *E. childrenae* only occur north of latitude 13ºN in Central and North America (Fig. 2a). We expanded the known distribution of the southern clade, notably filling gaps in northern South America, and concluded that it is distributed from Nicaragua to the Brazilian Amazon (Fig. 2a). We also extended and updated distribution records as follows: 1) *E. godartii* is distributed throughout the Pacific coast region, from Nicaragua to western Ecuador, including the Chocó-Darién and the Cauca river basin in Colombia (Fig. 2b); 2) *E. minyas* is found in the Magdalena valley and in the western Amazon Basin from Colombia to Mato Grosso, Brazil where it co-occurs with *E. toxana* 3) *E. toxana* occurs in the Amazon Basin from Obidos to the Andes and from Venezuela/Colombia to Mato Grosso, Brazil.

### Phylogenetic association between Eumaeus and Zamia

Eumaeus butterflies show remarkably strong host plant preferences (Fig. 3) with little evidence for overlap between species. Only *E. minyas* and *E. toxana*, which are sympatric in the Colombian Amazon, are likely to share host plants, although there are no explicit records in natural populations. The remaining species *E. childrenae, E. toxea*, and *E. godartii* do not overlap with other *Eumaeus* species in the host plants they use.

**Fig. 3.**
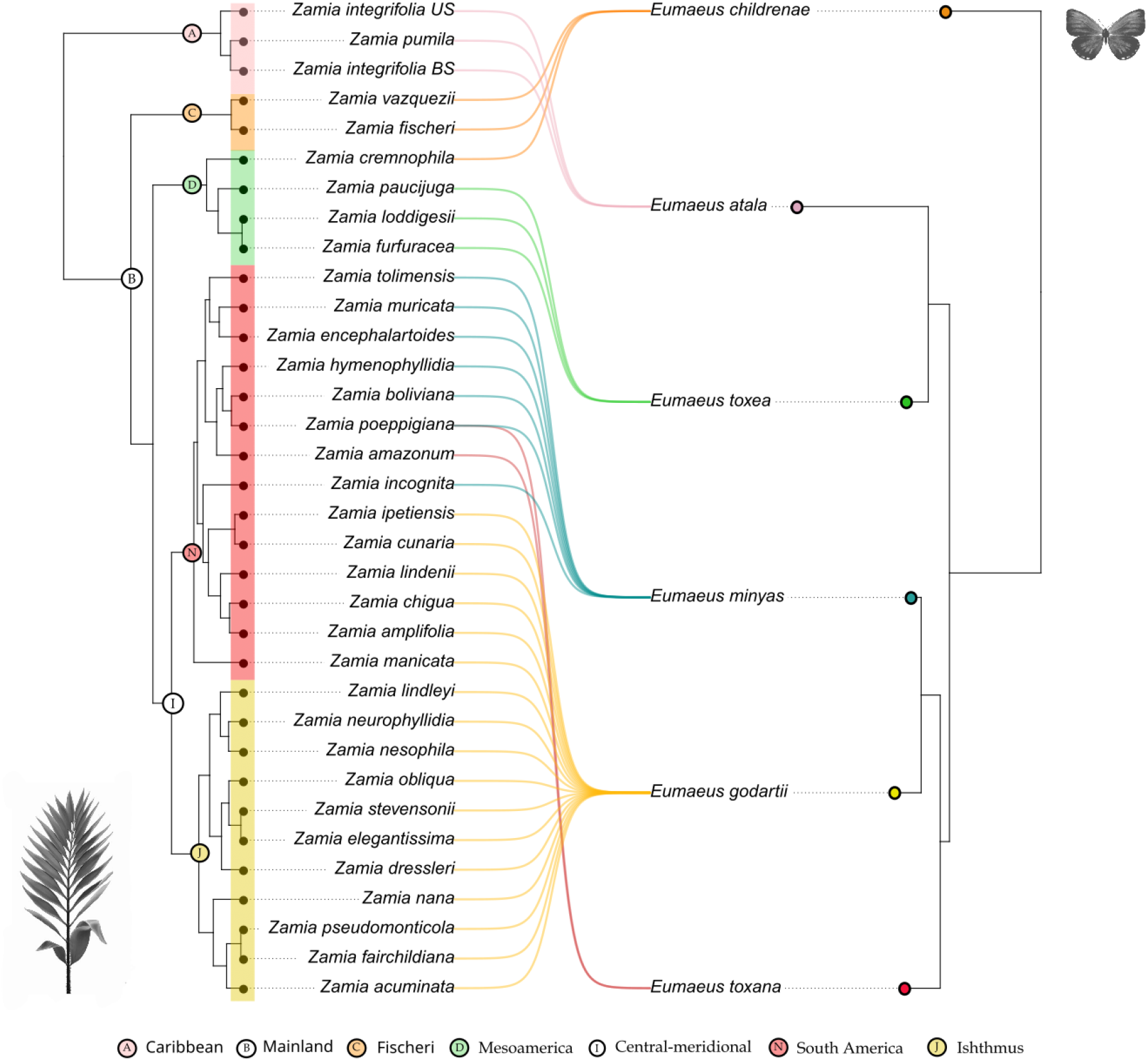
Phylogenetic association between *Eumaeus* and *Zamia* based on host plant records. *Zamia* phylogeny and *Zamia* clade names from Calonje et al. (2019).

The global-fit analysis revealed significant cophylogenetic signal between *Zamia* and *Eumaeus* (PACo: *m*^*2*^_*XY*_ =0.4488, *p* < 0.001, n = 1000, Fig. 4). Using a PGLMM, we found a significant effect of plant phylogeny (LRT x^2^ = 34.08, *p* < 0.001), but we did not find a similar effect of butterfly phylogeny (table 2). Thus, our results suggest that the phylogenetic signal is primarily driven by host plants, and this is congruent with the visual patterns of phylogenetic clustering apparent in Fig. 3.

**Table 2.**
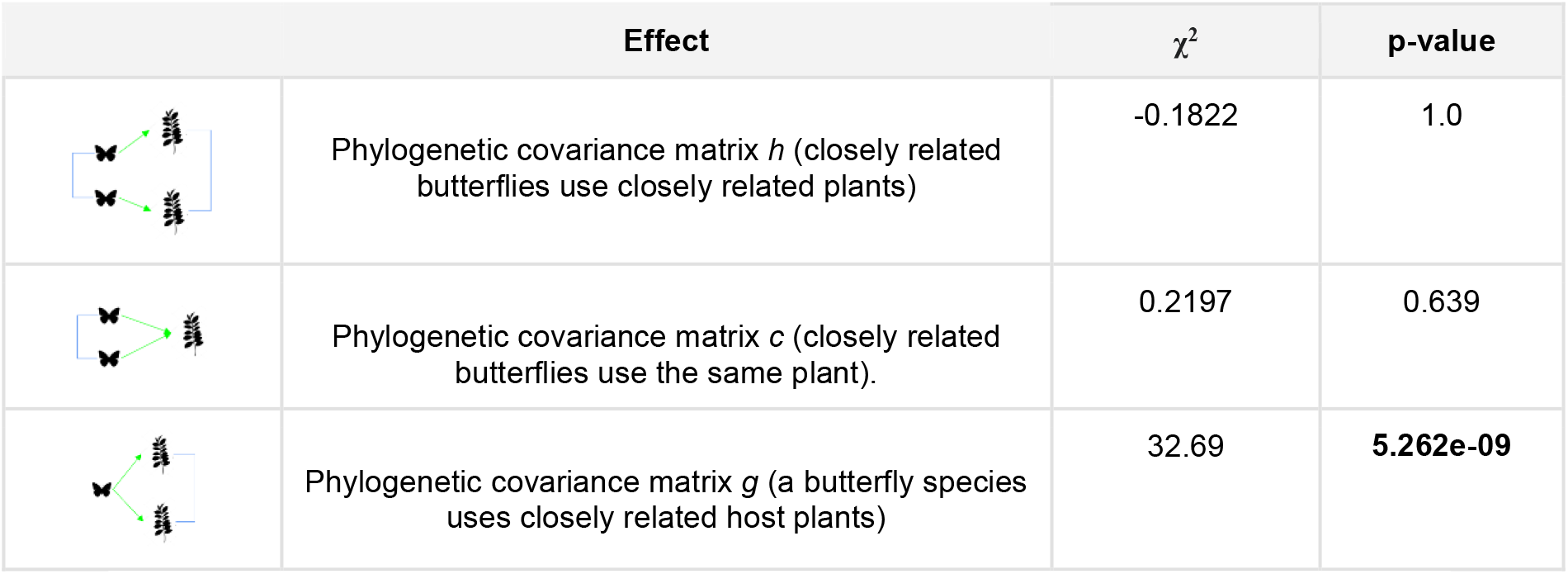
LRTs for the phylogenetic effects in the PGLMM of Zamia - Eumaeus interactions.

| | Effect | $\chi^2$ | p-value |
| --- | --- | --- | --- |
|  | Phylogenetic covariance matrix *h* (closely related butterflies use closely related plants) | -0.1822 | 1.0 |
|  | Phylogenetic covariance matrix *c* (closely related butterflies use the same plant). | 0.2197 | 0.639 |
|  | Phylogenetic covariance matrix *g* (a butterfly species uses closely related host plants) | 32.69 | **5.262e-09** |

**Fig. 4.**
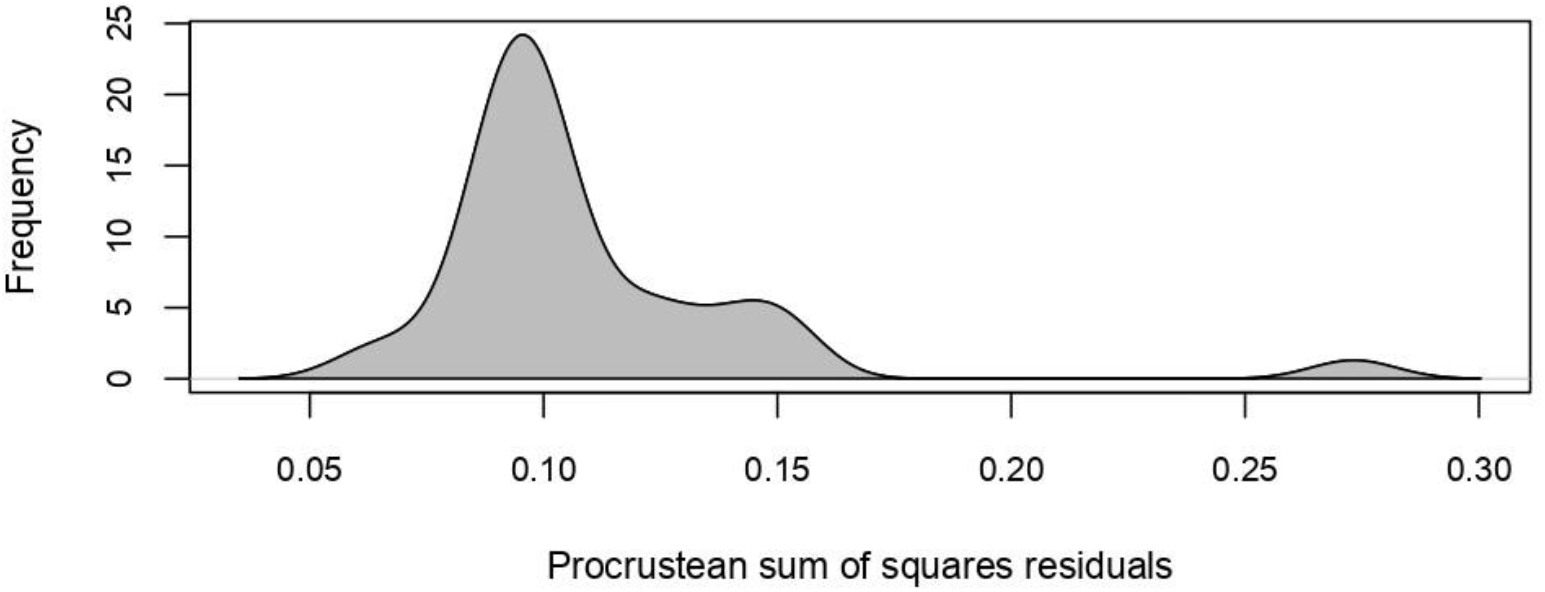
PACo permutation scores for the interaction matrix between species of *Eumaeus* and their *Zamia* host plants. The gray distribution depicts the permutation best fit Procrustean superimpositions. PACo returned statistically significant phylogenetic signal, implying that the interaction network between *Zamia* and *Eumaeus* is more congruent than permutated networks with randomized interactions links.

Unsurprisingly, the distributions of *Eumaeus* and *Zamia* are remarkably congruent (see Fig. 2a). Northerly distributed species of *Eumaeus* range from North America to Mexico and the Caribbean and are associated with *Zamia* species in the Caribbean and Mainland clades. *Eumaeus atala* uses solely the Caribbean species of *Zamia* (Fig. 3, A), although not all the zamias in this clade are restricted to the Caribbean. *E. childrenae* and *E. toxea* overlap in the Sierra Madre Forest and Mexican Drylands bioregion and use exclusively the Fisheri and Mesoamerica clades of *Zamia* (Fig. 3, C, D). Species of the *Eumaeus* southern clade use host plant species from the Central-meridional clade of *Zamia*, comprising the Isthmus and South America species (Fig. 3, I). *E*.*godartii* is the only species feeding on cycads from the Isthmus clade (Fig. 3, J), even though some species of host plant have ranges that extend to northern South America. Indeed, *E. godartii* caterpillars in South America were also recorded feeding only on species distributed West to the Andes, in the Pacific and Cunaria *Zamia* sub-clades (subclades following Calonje et. al., 2019). Contrastingly, *E. minyas* shares host plants in the Amazonian clade with *E. toxana*. Notably, no *Eumaeus* nor *Zamia* species are reported in the Guianas or Eastern Brazil.

### Divergence time in the Zamia and Eumaeus phylogenies

We found striking contemporaneity between *Zamia* and *Eumaeus* diversification (Fig. 5). Although *Zamia* evolution dates back to the Paleocene, the age of the last common ancestor of all extant species (mean age: 9.54 [9,10.64]) is remarkably similar to that of *Eumaeus* (mean age: 8.45 [6.11,10.47]) crown diversification in the Miocene. Notably, the most diverse clade of *Zamia*, the Mainland clade, evolved (mean age: 5.97 [3.92, 8.15]) synchronously with the core *Eumaeus* clade (mean age: 6.13 [4.33, 8.37]), which comprises all *Eumaeus* species except by *E. childrenae*. Similarly, the divergence of the Mesoamerica clade of *Zamia* (mean age: 3.9 [2.54, 5.34]) coincides with the split between Caribbean *E. atala* and mesoamerican *E. toxea* (mean age: 4.08 [2.12, 5.99]).

**Fig. 5.**
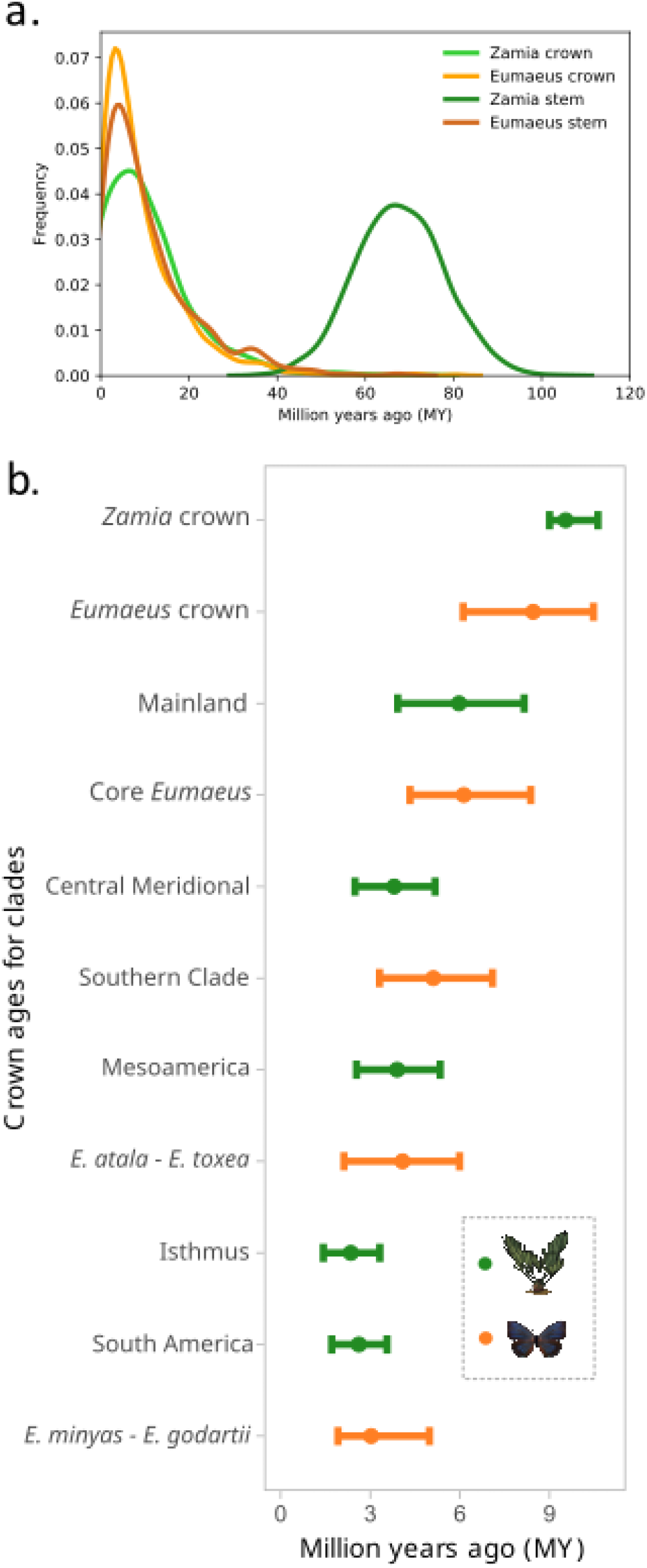
**a**. Divergence time estimation of *Zamia* and *Eumaeus* stem and crown groups with exponential priors. **b**. Mean and 95% confidence intervals for divergence times of the major *Zamia* and *Eumaeus* clades. All *Zamia* divergence times are from estimates in Calonje et al. (2019).

## Discussion

We present the first quantitative study of correlated phylogenies between interacting partners for a group of cycads and their herbivorous insects. Our analyses of interactions between *Eumaeus* butterflies and their host plant species of *Zamia* provide strong evidence of cophylogenetic signal between these clades. In addition, we found striking temporal overlap between the divergence times of the major *Zamia* and *Eumaeus* clades, further supporting a pattern of correlated evolution. We then dissected the pattern of phylogenetic congruence using mixed models and showed that the cophylogenetic signal is likely driven by phylogenetic tracking. This work represents a step towards a better understanding of the evolution of a recent yet specialized plant-herbivore interaction for one of the most ancient and endangered plant lineages.

Our cophylogenetic analysis shows a strong dependency between the evolutionary histories of the butterflies and their host plants, as predicted by Ehrlich and Raven’s (1964) arms race model of co-evolution. The latter posits that herbivory drives the evolution of novel defenses in plants, which then drives selection for traits to overcome them in butterflies. Each successful counteradaptation drives diversification, by placing the innovator in a new adaptive zone. Consequently, the interactions between plants and herbivores become phylogenetically structured over evolutionary time, with related herbivores feeding on closely related plants. To date, there have been no phylogenetic studies of the chemical defenses of cycads and their relationship with herbivores. However, there are many examples in angiosperms of toxic compounds with phylogenetic signal, such as cardenolides and glucosinolates in Brassicales (Züst et al., 2020) and *Asclepias* (Apocynaceae) or furanocoumarins in Umbelliferae (Berenbaum & Feeny, 1981). In these cases, adaptation to plant toxicity traits drives the specialization of beetle and butterfly herbivores, which accordingly show phylogenetically conserved host plant use (Berenbaum & Feeny, 1981; Rasmann & Agrawal, 2011; Wheat et al., 2007). It is therefore highly plausible that closely related *Zamia* species share similar chemical defenses that exert strong selective pressure for host conservatism in *Eumaeus*.

Nonetheless, it is important to note that congruent phylogenies of interacting taxa are neither necessary nor sufficient evidence of co-evolution, as they can also result from other processes (Blasco-Costa et al., 2021; Janzen, 1980; Russo et al., 2018). For example, congruent phylogenies might arise when both taxa are subject to the same vicariant events, which subdivide their populations and thereby result in correlated phylogenetic branching (Russo et al., 2018). Indeed, we observe parallel geographic structure in the phylogenies of both *Eumaeus* and *Zamia*. Calonje et al. (2019) showed that early divergent clades of *Zamia* are distributed in the Caribbean and Mesoamerica, with more recent and diverse clades distributed in southeastern Central Americas and South America. Similarly, we show here that early divergent species of *Eumaeus* range from the Caribbean to Mesoamerica, while the nested southern clade is restricted to the Isthmus and South America.

*Eumaeus godartii* is only found to the west of the Andes and uses the Isthmus clade of *Zamia* exclusively, suggesting the western cordillera of the Andes is a crucial geographic barrier. Similar phylogeographic breaks have also been observed in *Atta* ants, for which the Andes comprise an asymmetrical barrier to gene flow between populations in Central America and the rest of South America (Muñoz-Valencia et al., 2022). Likewise, clades in the hummingbird genus *Metallura* diverged and expanded from opposite sides of the Andes (Benham et al., 2015). However, the barrier effect of the Andes might not be as strong as previously thought for *Eumaeus* butterflies, or might have changed due to recent deforestation, since we found specimens exhibiting intermediate characteristics between *E. godartii* and *E. minyas* in the Magdalena and Cauca valleys. These regions may represent contact zones that allow gene-flow between these sister species, although they strongly diverge in their preferences for *Zamia* host plant species. Incomplete reproductive isolation between *E. minyas* and *E. godartii* is consistent with the short branch lengths subtending these species in the most recent whole-genome phylogeny (Robbins et al., 2021). Indeed, the Cauca valley is known to be a hybrid zone between other species of butterflies, such as *Heliconius* (Arias et al., 2008) and vertebrates, such as the Andean warbler (Céspedes-Arias et al., 2021).

Despite significant biogeographic correlations, the distributions and phylogeny of *Zamia* are not perfectly mirrored by *Eumaeus*. Notably, we observed biogeographic incongruences at deep nodes; the most ancient *Eumaeus* species are distributed in Mesoamerica, whereas the most ancient clade of *Zamia* diversified in the Caribbean. Furthermore, *E. childrenae* and *E. toxea* are not sister species, despite overlapping in the Sierra Madre Forest and the Mexican Drylands bioregion, where they use the Mesoamerican and Fisheri *Zamia* clades, respectively. Similarly, *E. toxana* and *E. minyas* are not sisters, although they co-occur throughout the Amazon Basin. This is in sharp contrast with *Zamia*, where there is well-known conservatism in their geographic ranges (i.e., closely related *Zamia* tend to inhabit the same geographic regions).

As well as vicariance, another process that can account for cophylogenetic signal without invoking co-evolution is phylogenetic tracking. Phylogenetic tracking is common when there are strong asymmetries in interaction strength, with one group relying more strongly upon the other (Russo et al., 2018). Under this scenario, parallel phylogenies result when the dependent species diverges and occupies niches created by speciation of the other (Russo et al., 2018). Thus, because one group tracks the speciation of the other group, diversification is usually asynchronous (Blasco-Costa et al., 2021; Ramírez et al., 2011). In the case of *Zamia* and *Eumaeus* interactions, host plants seem to exert stronger selection on their herbivores than the other way around. This asymmetry in selection is substantiated by previous work, showing that *Eumaeus* herbivory has little impact on cycad natural populations (Zabaleta Doria, 2013). Hence, it is likely that host plant use of *Eumaeus* tracks the evolution of their host plants. Nevertheless, we found remarkable temporal coincidence in plant and butterfly diversification (Fig. 5). In particular, we found that the stem age of *Eumaeus* is consistent with the crown age of extant *Zamia*. This result suggests rapid divergence of *Eumaeus* from its sister genus *Theorema* resulting from a host-shift from ancestral angiosperms hosts (Robbins et al., 2021), to the common ancestor of the most recent adaptive radiation of *Zamia* in the Miocene.

Qualitative observations about natural history and biogeography can generate useful hypotheses explaining cophylogenetic patterns observed between *Eumaeus* and *Zamia*. However, assessing the strength of association between the interaction matrix and the phylogenies offers a quantitative approach for disentangling co-evolution, vicariance, and phylogenetic tracking (Blasco-Costa et al., 2021; Russo et al., 2018). For instance, finding an effect of host plant phylogeny without an effect of butterfly phylogeny implies that phylogenetic signal is determined by the host plants, i.e., phylogenetic tracking (Blasco-Costa et al., 2021; Russo et al., 2018). In contrast, if the phylogenies of host plants and butterflies do not predict the interaction matrix, this implies that the branching patterns are similar because external selective forces such as vicariance are acting on both groups. Finally, the case that both phylogenies predict the interaction matrix is consistent with co-evolution, i.e., both partners exert selection on one other, which leads to reciprocal evolutionary change (Blasco-Costa et al., 2021; Russo et al., 2018).

To test these possibilities, we fit a mixed model with phylogenetic effects to the *Eumaeus-Zamia* interaction matrix. We found a highly significant effect of plant phylogeny, but not of butterfly phylogeny or the joint nested phylogenetic effect. Therefore, it is plausible that the significant global cophylogenetic signal found with `paco`, is primarily driven by the strong influence of the plant phylogeny predicting the bipartite network. As such, our results indicate that phylogenetic tracking is the most plausible mechanism underlying the significant cophylogenetic signal between *Zamia* and *Eumaeus*. This finding is also compatible with the original formulation of “sequential evolution”, in which plants drive the evolution of herbivorous insects, but without an appreciable selection feedback mechanism (Jermy, 1976).

Nonetheless, while phylogenetic tracking stands out as particularly relevant to this interaction network, we concur with Blasco-Costa et al. 2021 argument that vicariance, phylogenetic tracking, and co-evolution are not mutually exclusive. Patterns observed in ecological interactions are more plausibly a combination of processes, each acting with different intensity (Blasco-Costa et al., 2021). Particularly for an interaction exhibiting a strong level of host conservatism, such as the *Eumaeus-Zamia* (Calonje et al., 2010, 2011, 2015; López-Gallego, 2007; Segalla et al., 2021; Segalla & Calonje, 2019). In this study, we have consolidated an exhaustive dataset of host plant use by *Eumaeus* to start decoupling macroevolutionary patterns, but many gaps remain. We outline three main avenues for data acquisition to gain insight into the extent of codivergence between cycads and *Eumaeus* herbivores.

First, collecting data on interaction traits, such as frequency of visits, abundance, number of eggs laid, leaf damage, or sensory biases, can provide a more comprehensive picture of the eco-evolutionary dynamics of this two-sided interaction. For instance, the characterization of host defensive traits across the *Zamia* phylogeny can help to explain the strong phylogenetic signal on host plant use by *Eumaeus* butterflies. Second, while genomic resources are now available for all *Eumaeus* species (Robbins et al., 2021), next-generation sequencing of cycads has lagged behind. Genomes, transcriptomes, and metabolomes of *Zamia* species would enable profiling clade-specific chemical defenses, which could in turn be linked to *Eumaeus* counterdefenses. Third, our approach of scoring presence-absence of interactions based solely on morphology and distribution might not properly account for cryptic diversity. Since the greatest diversity of *Zamia* species is found in South America, we anticipate finding structured populations in the southern *Eumaeus* clade concomitant with the extensive diversity of this host plant group in this continent. Cryptic diversity might be especially prevalent in aposematic butterflies such as *Eumaeus* whereby positive frequency-dependent selection can constrain the phenotypic divergence of traits traditionally used in Lepidoptera taxonomy, such as wing color patterns. Exhaustive genomic sampling at the population level across broad geographical scales is therefore necessary to elucidate the extraordinary recent evolution of this ancient plant lineage and their unique and specialized herbivores.

## Acknowledgements

We thank the Colombian Cycad Society for funding and logistical support for this project. We also thank the Xerces Society for Invertebrate Conservation for a DeWind award to WAVM to study endangered *Eumaeus* butterflies. We are grateful towards the Montgomery Botanical Center (MBC) for the unconditional support of cycad research. We thank Jonathan Castro, whose previous work provided us with valuable data about *Eumaeus* host plants in Colombia and to Cornelio Bota, who allowed us to use one of his photographs. We are also grateful towards Jing Zhang and Nick Grishin for sharing sequencing data of *Eumaeus* and allies. We thank all of the curators and researchers at museums in Colombia, specially Juliana Cardona-Duque and Carolina Vélez at CBUCES, Martha Wolff at CEUA, Gonzalo Andrade at the ICN, John Cesar Neita and Johann Cardenas at IaVH for their kind disposition in the collections. We are especially grateful to our field assistants in Colombian forests: the Rivera Mejía family, Miguel González, Rosendo Yukuna Matapí, and Miguel Arcángel Bora for their knowledge and guidance through their territories. We are also grateful to Luis Horacio Agudelo and Vanessa Correa for their help during fieldwork and the Botero-Sierra family for their unconditional support.

## References

Arias, C. F., Muñoz, A. G., Jiggins, C. D., Mavárez, J., Bermingham, E., & Linares, M. (2008). A hybrid zone provides evidence for incipient ecological speciation in Heliconius butterflies. Molecular Ecology, 17(21), 4699–4712. https://doi.org/10.1111/j.1365-294X.2008.03934.x

Balbuena, J. A., Míguez-Lozano, R., & Blasco-Costa, I. (2013). PACo: A Novel Procrustes Application to Cophylogenetic Analysis. PLOS ONE, 8(4), e61048. https://doi.org/10.1371/journal.pone.0061048

Benham, P. M., Cuervo, A. M., McGuire, J. A., & Witt, C. C. (2015). Biogeography of the Andean metaltail hummingbirds: Contrasting evolutionary histories of tree line and habitat-generalist clades. Journal of Biogeography, 42(4), 763–777. https://doi.org/10.1111/jbi.12452

Berenbaum, M., & Feeny, P. (1981). Toxicity of Angular Furanocoumarins to Swallowtail Butterflies: Escalation in a Coevolutionary Arms Race? Science, 212(4497), 927– 929. https://doi.org/10.1126/science.212.4497.927

Blasco-Costa, I., Hayward, A., Poulin, R., & Balbuena, J. A. (2021). Next-generation cophylogeny: Unravelling eco-evolutionary processes. Trends in Ecology & Evolution, 36(10), 907–918. https://doi.org/10.1016/j.tree.2021.06.006

Bowers, M. D., & Larin, Z. (1989). Acquired chemical defense in the lycaenid butterfly, Eumaeus atala. Journal of Chemical Ecology, 15(4), 1133–1146. https://doi.org/10.1007/BF01014817

Calonje, M., Esquivel, H. E., Stevenson, D., Calonje, C., & Pava, D. (2011). A new arborescent species of Zamia from the Central Cordillera of Tolima, Colombia (Cycadales, Zamiaceae), with comments on the Z. poeppigiana species complex. Brittonia, 63(4), 442–451. https://doi.org/10.1007/s12228-011-9190-4

Calonje, M., Morales, G., López-Gallego, C., & Roldán, F. J. (2015). A taxonomic revision of Zamia montana and Zamia oligodonta, with notes on their conservation status. Phytotaxa, 192(4), 279. https://doi.org/10.11646/phytotaxa.192.4.5

Calonje, M., Stevenson, D., Calonje, C., Ramos, Y. A., & Lindstrom, A. (2010). A new species of Zamia from Chocó, Colombia (Cycadales, Zamiaceae). Brittonia, 62(1), 80–85. https://doi.org/10.1007/s12228-009-9094-8

Castillo-Guevara, C., & Rico-Gray, V. (2002). Is cycasin in Eumaeus minyas (Lepidoptera: Lycaenidae) a predator deterrent? Interciencia, 27(9), 465–470.

Castillo-Guevara, C., & Rico-Gray, V. (2003). The role of macrozamin and cycasin in cycads (Cycadales) as antiherbivore defenses. Journal of the Torrey Botanical Society, 130(3), 206–217. https://doi.org/10.2307/3557555

Céspedes-Arias, L. N., Cuervo, A. M., Bonaccorso, E., Castro-Farias, M., Mendoza-Santacruz, A., Pérez-Emán, J. L., Witt, C. C., & Cadena, C. D. (2021). Extensive hybridization between two Andean warbler species with shallow divergence in mtDNA. Ornithology, 138(1), 1–28. https://doi.org/10.1093/ornithology/ukaa065

Charlton, T. S., Marini, A. M., Markey, S. P., Norstog, K., & Duncan, M. W. (1992). Quantification of the neurotoxin 2-amino-3-(methylamino)-propanoic acid (BMAA) in cycadales. Phytochemistry, 31(10), 3429–3432. https://doi.org/10.1016/0031-9422(92)83700-9

Clark, D. B., & Clark, D. A. (1991). Herbivores, herbivory, and plant phenology: Patterns and consequences in a tropical rain-forest cycad. In Plant-animal interactions: Evolutionary ecology in tropical and temperate regions (Wiley-Interscience, p. 639). Wiley.

Contreras-Medina, R., Ruiz-Jiménez, C. A., & Luna Vega, I. (2003). Caterpillars of Eumaeus childrenae (Lepidoptera: Lycaenidae) feeding on two species of cycads (Zamiaceae) in the Huasteca region, Mexico. Revista De Biologia Tropical, 51(1), 201–203.

De Luca, P., Moretti, A., Sabato, S., & Siniscalco Gigliano, G. (1980). The ubiquity of cycasin in cycads. Phytochemistry, 19(10), 2230–2231. https://doi.org/10.1016/S0031-9422(00)82239-6

Duncan, M. W., Kopin, I. J., Garruto, R. M., Lavine, L., & Markey, S. P. (1988). 2-Amino-3 (methylamino)-propionic acid in cycad-derived foods is an unlikely cause of amyotrophic lateral sclerosis/parkinsonism. Lancet (London, England), 2(8611), 631– 632. https://doi.org/10.1016/s0140-6736(88)90671-x

Espeland, M., Breinholt, J., Willmott, K. R., Warren, A. D., Vila, R., Toussaint, E. F. A., Maunsell, S. C., Aduse-Poku, K., Talavera, G., Eastwood, R., Jarzyna, M. A., Guralnick, R., Lohman, D. J., Pierce, N. E., & Kawahara, A. Y. (2018). A Comprehensive and Dated Phylogenomic Analysis of Butterflies. Current Biology, 28(5), 770-778.e5. https://doi.org/10.1016/j.cub.2018.01.061

Fuzessy, L., Silveira, F. A. O., Culot, L., Jordano, P., & Verdú, M. (2022). Phylogenetic congruence between Neotropical primates and plants is driven by frugivory. Ecology Letters, 25(2), 320–329. https://doi.org/10.1111/ele.13918

González, F. (2004). De Colombia, Zamia encephalartoides (Zamiaceae) Por Parte De Eumaeus (Lepidoptera: Lycaenidae). Revista de La Academia Colombiana de Ciencias Exactas, Fisicas y Naturales, XXVIII, 233–244.

Goodson, F. W. (1947). Notes on the genus Eumaeus Hübner (Lep., Lycaenidae). The Entomologist, 80, 273–276.

Hutchinson, M. C., Cagua, E. F., Balbuena, J. A., Stouffer, D. B., & Poisot, T. (2017). paco: Implementing Procrustean Approach to Cophylogeny in R. Methods in Ecology and Evolution, 8(8), 932–940. https://doi.org/10.1111/2041-210X.12736

IUCN Red List of Threatened Species. (2022). https://www.iucnredlist.org/resources/summary-statistics

Janzen, D. H. (1980). When Is It Coevolution? Evolution, 34(3), 611–612. https://doi.org/10.1111/j.1558-5646.1980.tb04849.x

Jiménez-Pérez, N. del C., Moguel-Ordóñez, E. J., Hernández-Jiménez, O. A., & Cruz, M. P. la. (2017). Primer Registro de Eumaeus childrenae Sobre la Cícada Microendémica Zamia cremnophila (Zamiaceae) en Tabasco, México. Southwestern Entomologist, 42(2), 609–612. https://doi.org/10.3958/059.042.0233

Koi, S., & Daniels, J. (2015). New and revised life history of the Florida hairstreak Eumaeus atala (Lepidoptera: Lycaenidae) with notes on its current conservation status. Florida Entomologist, 98(4), 1134–1147. https://doi.org/10.1653/024.098.0418

Koi, S., & Daniels, J. (2017). Life History Variations and Seasonal Polyphenism in Eumaeus atala (Lepidoptera: Lycaenidae). Florida Entomologist, 100(2), 219–229. https://doi.org/10.1653/024.100.0216

Koi, S., & Hall, D. W. (2016). Atala Butterfly, Atala Hairstreak, Coontie Hairstreak, Eumaeus atala Poey 1832 (Insecta: Lepidoptera: Lycaenidae). Extension Document of the Entomology and Nematology Department, UF/IFAS Extension, 1832(EENY-641), 1– 11.

Laqueur, G. L., & Spatz, M. (1968). Toxicology of cycasin. Cancer Research, 28(11), 2262– 2267.

Li, D., Dinnage, R., Nell, L. A., Helmus, M. R., & Ives, A. R. (2020). phyr: An r package for phylogenetic species-distribution modelling in ecological communities. Methods in Ecology and Evolution, 11(11), 1455–1463. https://doi.org/10.1111/2041-210X.13471

López-Gallego, C. (2007). Effects of habitat degradation on the evolutionary dynamics of populations in a rainforest cycad (Gymnospermae) [University of New Orleans]. https://www.proquest.com/openview/986321eb565c362bd4ec7f5d5de177e9/1?cbl=18750&pq-origsite=gscholar

López-Gallego, C., Olaya-Rodríguez, M. H., Velásquez-Tibata, J., & Noguera-Urbano, E. (2019). Atlas de la biodiversidad de Colombia. Zamias. Instituto de Investigación de Recursos Biológicos Alexander von Humboldt.

Martínez-Lendech, N., Córdoba-Aguilar, A., & Serrano-Meneses, M. A. (2007). Body size and fat reserves as possible predictors of male territorial status and contest outcome in the butterfly Eumaeus toxea Godart (Lepidoptera: Lycaenidae). Journal of Ethology, 25(2), 195–199. https://doi.org/10.1007/s10164-007-0040-5

Muñoz-Valencia, V., Vélez-Martínez, G. A., Montoya-Lerma, J., & Díaz, F. (2022). Role of the Andean uplift as an asymmetrical barrier to gene flow in the neotropical leaf-cutting ant Atta cephalotes. Biotropica, 54(1), 191–204. https://doi.org/10.1111/btp.13050

Nagalingum, N. S., Marshall, C. R., Quental, T. B., Rai, H. S., Little, D. P., & Mathews, S. (2011). Recent Synchronous Radiation of a Living Fossil. Science, 334(6057), 796– 799. https://doi.org/10.1126/science.1209926

Norstog, K. J., & Fawcett, P. K. S. (1989). Insect-Cycad Symbiosis and Its Relation to the Pollination of Zamia furfuracea (Zamiaceae) by Rhopalotria mollis (Curculionidae). American Journal of Botany, 76(9), 1380–1394. https://doi.org/10.1002/j.1537-2197.1989.tb15117.x

Norstog, K., & Nicholls, T. J. (1997). The Biology of the Cycads. Comstock Pub. Associates.

Pierce, N. E., Braby, M. F., Heath, A., Lohman, D. J., Mathew, J., Rand, D. B., & Travassos, M. A. (2002a). The Ecology and Evolution of Ant Association in the Lycaenidae (Lepidoptera). Annual Review of Entomology, 47(1), 733–771. https://doi.org/10.1146/annurev.ento.47.091201.145257

QGIS.org, %Y. QGIS Geographic Information System. QGIS Association. http://www.qgis.org

Rafferty, N. E., & Ives, A. R. (2013). Phylogenetic trait-based analyses of ecological networks. Ecology, 94(10), 2321–2333. https://doi.org/10.1890/12-1948.1

Ramírez, S. R., Eltz, T., Fujiwara, M. K., Gerlach, G., Goldman-Huertas, B., Tsutsui, N. D., & Pierce, N. E. (2011). Asynchronous Diversification in a Specialized Plant-Pollinator Mutualism. Science, 333(6050), 1742–1746. https://doi.org/10.1126/science.1209175

Rasmann, S., & Agrawal, A. A. (2011). Evolution of Specialization: A Phylogenetic Study of Host Range in the Red Milkweed Beetle (Tetraopes tetraophthalmus). The American Naturalist, 177(6), 728–737. https://doi.org/10.1086/659948

Robbins, R. K., Cong, Q., Zhang, J., Shen, J., Riera, J. Q., Murray, D., Busby, R. C., Faynel, C., Hallwachs, W., Janzen, D. H., & Grishin, N. V. (2021). A switch to feeding on cycads generates parallel accelerated evolution of toxin tolerance in two clades of Eumaeus caterpillars (Lepidoptera: Lycaenidae). Proceedings of the National Academy of Sciences of the United States of America, 118(7). https://doi.org/10.1073/pnas.2018965118

Rothschild, M., Nash, R. J., & Bell, E. A. (1986). Cycasin in the endangered butterfly Eumaeus atala florida. Phytochemistry, 25(8), 1853–1854. https://doi.org/10.1016/S0031-9422(00)81161-9

Ruiz–García, N. (2020). Effectiveness of the aposematic Eumaeus childrenae caterpillars against invertebrate predators under field conditions. Animal Biodiversity and Conservation, 43(1), 109–114. https://doi.org/10.32800/abc.2020.43.0109

Russo, L., Miller, A. D., Tooker, J., Bjornstad, O. N., & Shea, K. (2018). Quantitative evolutionary patterns in bipartite networks: Vicariance, phylogenetic tracking or diffuse co-evolution? Methods in Ecology and Evolution, 9(3), 761–772. https://doi.org/10.1111/2041-210X.12914

Salzman, S., Whitaker, M., & Pierce, N. E. (2018). Cycad-feeding insects share a core gut microbiome. Biological Journal of the Linnean Society, 123(4), 728–738. https://doi.org/10.1093/biolinnean/bly017

Santos Murgas, A., & Abrego, J. C. (2016). Historia Natural de Eumaeus godartii (Lycaenidae, Lepidoptera) y Herbivoría en Zamia manicata. Revista Colón Ciencias, Tecnologia y Negocios, 3(1), 36–48.

Schneider, D., Wink, M., Sporer, F., & Lounibos, P. (2002). Cycads: Their evolution, toxins, herbivores and insect pollinators. Naturwissenschaften, 89(7), 281–294. https://doi.org/10.1007/s00114-002-0330-2

Segalla, R., & Calonje, M. (2019). Zamia brasiliensis, a new species of Zamia (Zamiaceae, Cycadales) from Mato Grosso and Rondônia, Brazil. Phytotaxa, 404(1), 1. https://doi.org/10.11646/phytotaxa.404.1.1

Segalla, R., Pinheiro, F., & Morellato, L. P. C. (2021). Reproductive biology of the South American cycad Zamia boliviana, involving brood-site pollination. Plant Species Biology, 36(2), 348–360. https://doi.org/10.1111/1442-1984.12322

Tang, W. (1987). Heat Production in Cycad Cones. Botanical Gazette, 148(2), 165–174. https://doi.org/10.1086/337644

Taylor B. A. S. (2020). Eumaeus godartii butterfly: Pest friend or foe? In C. Lopez-Gallego, M. Calonje, M. Griffith, & J. Khuraijam, Proceedings of Cycad 2015: The 10th International Conference on Cycad Biology.

Toon, A., Terry, L. I., Tang, W., Walter, G. H., & Cook, L. G. (2020). Insect pollination of cycads. Austral Ecology, 45(8), 1033–1058. https://doi.org/10.1111/AEC.12925

Valencia-Montoya, W. A., Quental, T. B., Tonini, J. F. R., Talavera, G., Crall, J. D., Lamas, G., Busby, R. C., Carvalho, A. P. S., Morais, A. B., Oliveira Mega, N., Romanowski, H. P., Liénard, M. A., Salzman, S., Whitaker, M. R. L., Kawahara, A. Y., Lohman, D. J., Robbins, R. K., & Pierce, N. E. (2021). Evolutionary trade-offs between male secondary sexual traits revealed by a phylogeny of the hyperdiverse tribe Eumaeini (Lepidoptera: Lycaenidae). Proceedings of the Royal Society B: Biological Sciences, 288(1950). https://doi.org/10.1098/rspb.2020.2512

Valencia-Montoya, W. A., Tuberquia, D., Guzmán, P. A., & Cardona-Duque, J. (2017). Pollination of the cycad Zamia incognita A. Lindstr. & Idárraga by Pharaxonotha beetles in the Magdalena Medio Valley, Colombia: A mutualism dependent on a specific pollinator and its significance for conservation. Arthropod-Plant Interactions, 11(5), 717–729. https://doi.org/10.1007/s11829-017-9511-y

Vega, A., & Bell, E. A. (1967). α-Amino-β-methylaminopropionic acid, a new amino acid from seeds of Cycas circinalis. Phytochemistry, 6(5), 759–762. https://doi.org/10.1016/S0031-9422(00)86018-5

Warren, A. D., Davis, K. J., & Stangeland, E. M. (2016). Lycaenidae of the Americas 21-XI-2017. Buttterflies of America. https://www.butterfliesofamerica.com/L/Lycaenidae.htm

Wheat, C. W., Vogel, H., Wittstock, U., Braby, M. F., Underwood, D., & Mitchell-Olds, T. (2007). The genetic basis of a plant–insect coevolutionary key innovation. Proceedings of the National Academy of Sciences, 104(51), 20427–20431. https://doi.org/10.1073/pnas.0706229104

Whitaker, M. R. L., & Salzman, S. (2020). Ecology and evolution of cycad-feeding Lepidoptera. Ecology Letters, 23(12), 1862–1877. https://doi.org/10.1111/ele.13581

Zabaleta Doria, D. (2013). Patrones de herbivoría en poblaciones de cycadas de diferentes hábitats y para distintos tipos de herbivoros. [Universidad de Antioquia].

Zhifeng, G., & Thomas, B. A. (1989). A review of fossil cycad megasporophylls, with new evidence of Crossozamia pomel and its associated leaves from the lower permian of Taiyuan, China. Review of Palaeobotany and Palynology, 60(3), 205–223. https://doi.org/10.1016/0034-6667(89)90044-4

Züst, T., Strickler, S. R., Powell, A. F., Mabry, M. E., An, H., Mirzaei, M., York, T., Holland, C. K., Kumar, P., Erb, M., Petschenka, G., Gómez, J.-M., Perfectti, F., Müller, C., Pires, J. C., Mueller, L. A., & Jander, G. (2020). Independent evolution of ancestral and novel defenses in a genus of toxic plants (Erysimum, Brassicaceae). ELife, 9, e51712. https://doi.org/10.7554/eLife.51712

